# Using Biogeography to Assess Key Adaptation Strength in Two Bird Families

**DOI:** 10.1101/214148

**Authors:** Abdel H. Halloway, Christopher J. Whelan, Çağan H. Şekercioğlu, Joel S. Brown

## Abstract

Adaptations can be thought of as evolutionary technologies which allow an organism to exploit environments. Among convergent taxa, adaptations may be largely equivalent with the taxa operating in a similar set of environmental conditions, divergent with the taxa operating in different sets of environmental conditions, or superior with one taxon operating within an extended range of environmental conditions than the other. With this framework in mind, we sought to characterize the adaptations of two convergent nectarivorous bird families, the New World hummingbirds (Trochilidae) and Old World sunbirds (Nectariniidae), by comparing their biogeography. Looking at their elevational and latitudinal gradients, hummingbirds not only extend into but also maintain species richness in more extreme environments. We suspect that hummingbirds have a superior key adaptation that sunbirds lack, namely a musculoskeletal architecture that allows for hovering. Through biogeographic comparisons, we have been able to assess and understand adaptations as evolutionary technologies among two convergent bird families, a process that should work for most taxa.

## Introduction

Convergent evolution provides startling examples of how natural selection shapes the traits of species to optimize fitness (1). Species’ morphologies, physiologies, and behaviors become fine-tuned to their shared ecologies somewhat irrespective of evolutionary history (2). Besides the species level, convergence can happen between higher taxa (e.g., families and orders). One striking example is the convergence between the New World nectarivorous hummingbirds (order Apodiformes, family Trochilidae) and the Old World passerine nectarivores including the Hawaiian honeycreepers (order Passeriformes, family Fringillidae), Australian honeyeaters (order Passeriformes, family Meliphagidae) and the Asian and African sunbirds (order Passeriformes, family Nectariniidae). Some or all members of these families show convergent adaptations for nectarivory, particularly elongated bills and extensile tongues. Their remarkable convergence, especially between hummingbirds and sunbirds, makes them ripe for analysis of adaptations as evolutionary technologies.

Adaptations can be thought of as evolutionary technologies that allow an organism to operate within an environment. Among evolutionary convergent taxa, adaptations might be equivalent leading to similar fitness in similar environmental conditions, e.g. the convergent snake constrictor families Boidae and Pythonidae (Fig. 1a) (3). In this case, both clades operate under similar fundamental niches. Differences in adaptations, though, can change the fundamental niches of the convergent clades and open new ecological opportunities (4). Such adaptations are known as as key adaptations. Key adaptations may be divergent evolutionary technologies with the taxa occupying different fundamental niches, e.g. the ankle bones of the grandorder Euarchonta (four orders of mammals including primates) that promote arboreal living (Fig. 1b). Hummingbirds and hawkmoths (order Lepidoptera, family Sphingidae) are instructive examples of convergent families displaying divergent evolutionary technologies. As vertebrates and invertebrates respectively, they strongly differ in virtually all aspects of ontogeny, morphology, and physiology. Yet, their coexistence in the New World suggests that one set of evolutionary technologies is not superior to the other under all circumstances leading to a partitioning of environmental conditions (5).

**Fig. 1.**
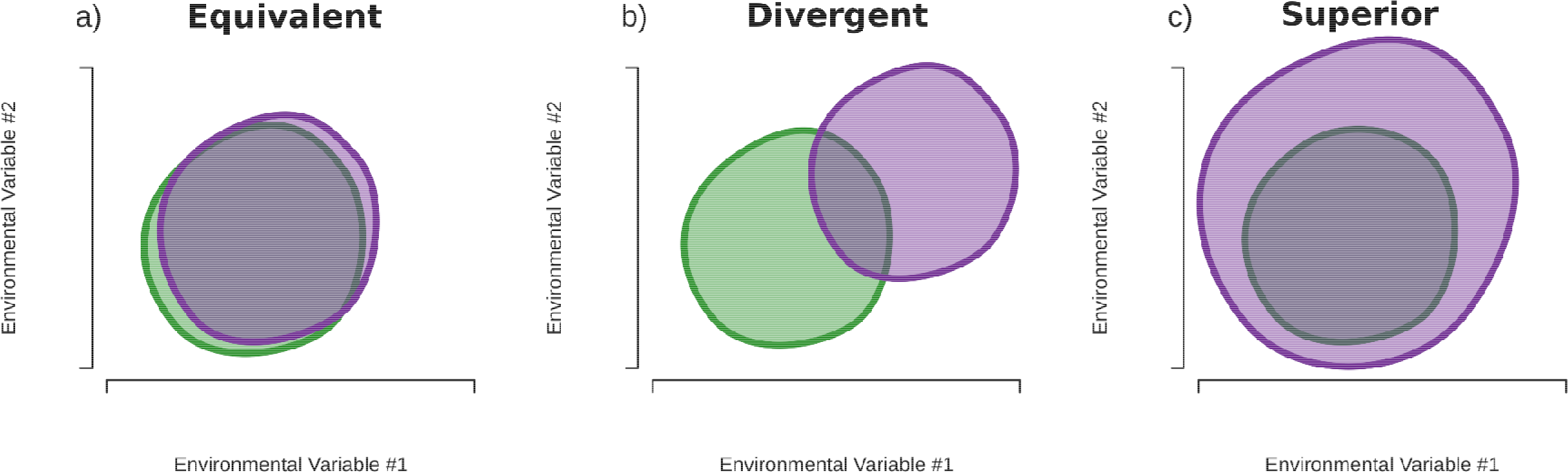
How differences in adaptations between two convergent taxa (purple and green) may lead may alter the set of environmental variables (its fundamental niche) under which a taxon has positive fitness (colored region). (a) Adaptations are largely equivalent, and the taxa survive under similar conditions. (b) Adaptations are divergent, and there is little overlap in environmental conditions in which both taxa survive. (c) The superior adaptations of the purple clade mean it can survive under a greater set of environmental variables (i.e., it has a larger fundamental niche) compared to the green clade.

Key adaptations may also be superior evolutionary technologies allowing a taxon to expand beyond its original range of environments and have a greater fundamental niche (Fig. 1c). Examples of superior key adaptations include the retractable necks among turtles of the suborder Cryptodira, which protect them from predation, and the infrared-sensing pits among vipers of the subfamily Crotalinae, which allows them to “see” mammals at night. When a key adaptation creates a superior evolutionary technology, we might see the replacement of another clade – typically but not always the ancestral clade – by the new one (6; 7; 8). Or the ancestral clade may persist where the derived clade has yet to colonize as is the case with the Pleurodira turtles of the Southern hemisphere. Since hummingbirds and sunbirds do not occur sympatrically and therefore do not interact, it is hard to discern whether they represent equivalent, divergent, or superior evolutionary technologies. We hypothesize that hummingbirds represent the latter compared to sunbirds and other nectarivorous passerines with hummingbirds possessing a superior key adaptation making them an example of “progressive evolution” (9; 6).

Hummingbirds display a stronger mutualistic co-adaptation with flowers compared to sunbirds (10; 11). All hummingbirds feed almost exclusively on nectar, only supplementing protein intake by eating small insects (12). As such, they have evolved distinct anatomical and morphological features suited to nectar foraging. In addition to an elongated bill and extensile tongue, the hummingbird’s tongue acts as a micro-pump for reaching and gathering nectar (13; 14). They possess large breast muscles (30% of body weight), skeletal architecture common to Apodiformes, and dense erythrocyte counts for delivering a steady supply of oxygen to feed extremely active muscles (10). Specialized wings allow hummingbirds to hover and fly backwards. Sunbirds, on the other hand, are not as tightly adapted to nectar feeding with many species supplementing their diet with insects, seeds, fruit, and flower heads, and others being largely insectivorous (15). They also show large variation in bill and flight morphology with the flowerpeckers and the *Hedydipna* and *Hypogramma* sunbirds having broad, flat tongues. In addition, all sunbirds lack the musculoskeletal architecture to hover and must perch to feed (11). These anatomical differences along with differences in species richness (364 hummingbird species vs. 147 sunbird species) suggest that hummingbirds have a superior key adaptation not found in sunbirds (16). Furthermore, the geographic isolation between the taxa allowed for independent diversification, making them ideal convergent clades to assess adaptations.

Testing for a key adaptation requires two things: elucidating a mechanistic hypothesis for its ecological and functional role and a comparison between clades (7). When comparing clades, species richness and diversification rates have typically been used (4). Besides these properties, we also surmise that a greater geographical range would be seen with a superior key adaptation. By increasing net fitness overall, a superior key adaptation should increase the fitness of a clade at the margins of its range (17); therefore, clades with a superior key adaptation will be better able to handle abiotic stress and live under harsher climatic regimes. Looking at the convergent mice genera *Peromyscus* and *Apodemus*, *Peromyscus* inhabits colder, more arid, and higher habitats compared to *Apodemus* due to its more efficient and widely used torpor state (18; 19; 20). Between species richness and biogeography, comparing the latter may be more useful to assess adaptations as evolutionary technologies since biogeographical extent explicitly depends upon a clades’ overall net fitness and directly tests its ecological role.

To characterize the adaptations of hummingbirds and sunbirds, we compared their biogeography by analyzing each family’s latitudinal and elevational distribution. We demonstrate that hummingbirds as a clade inhabit more extreme latitudes and maintain their species richness at higher elevations. We hypothesize these differences in biogeography reveal a superior key adaptation present in hummingbirds but absent in sunbirds. We speculate that the key adaptation may be either the unique tongue of the hummingbirds or the unique wing architecture that allows for hovering. We further speculate on the role adaptations as evolutionary technologies play in influencing an organism’s ability to exploit the environment.

## Materials and Methods

To assess the adaptations of hummingbirds and sunbirds, we gathered each family’s latitudinal and elevational gradient of species richness. These gradients are robust geographic patterns that generally show species richness declining towards higher altitudes and more extreme latitudes (21; 22; 23; 24; 25; 26; 27). Numerous environmental properties change along both gradients. Aridity declines significantly around 30 to 40 degrees latitude; a thinner atmosphere and more variable daily temperatures occur with increased elevation; and more variable seasonal temperatures, less productivity, and colder temperatures occur with both. Taxa with superior evolutionary technologies should be better able to deal with these challenges (28).

To compare the biogeography of hummingbirds and sunbirds, we gathered the latitudinal and elevational range of all species from each family. Elevational ranges came from a global bird ecology database covering all the bird species of the world (29) while latitudinal ranges of the families were taken from shapefiles downloaded from BirdLife International and NatureServe with data extracted using R packages “sp”, “raster”, “rasterVis”, “maptools”, and “rgeos” (30). All latitudinal extremes located in the Southern hemisphere were converted to negative values, and latitudinal maxima and minima were rounded up and down to the nearest integer respectively. For example, the hummingbird species *Amazilia amabilis* which ranges from 14.17N to 3.98S would have its range taken as 15 to −4. An additional measure of distance from the equator, hereafter referred to as “polewardness”, was created. If a species’ range crossed the equator, then the poleward range was taken to be from 0 to the maximum distance from the equator. For *A. amabilis*, its poleward range would be 0 to 15 degrees. The poleward range of an only Northern or Southern hemispheric species would simply be the absolute value of its latitudinal range.

With ranges in hand, we compared the families in two ways. First, we compared several empirical cumulative distribution functions (ECDFs) based upon the three geographical properties (elevation, latitude, and polewardness) for each family. Each ECDF started from sea level, the South Pole, and the equator and traced to higher altitudes, northward, and more extreme poles. For each geographical property, three ECDFs were created with a species’ presence based on the minimum, the midpoint, and the maximum of its range. Since species which cross the equator are not necessarily symmetric about it, the midpoint of a species poleward range may not accurately reflect its bias towards the equator or poles. Therefore, we created another measurement of species presence for polewardness, its expected value (see SI). This led to ten different ECDFs for each family: minimum, maximum, and midpoint for elevation, latitude, and polewardness and the additional measure of expected polewardness. Each type of ECDF was then compared between families using the Kolmogorov-Smirnov and Anderson-Darling minimum difference estimation (MDE) tests with the assumption that the hummingbird ECDF is less than the sunbird ECDF (one-tailed tests).

The ECDF analysis tells us whether the distributions differ, not necessarily how they differ. Therefore, we additionally sought to characterize each family’s distribution by measuring changes in species richness with polewardness and elevation. To do so, we first counted the number of species in poleward and elevation intervals of 5 degrees and 500 meters for each family. If the edge of a species’ range was at the cutoff point of the interval, it would be considered present in the lower interval but not in the upper interval due to previous rounding. In the example with *A. amabilis*, this would mean that the species is counted in the 10 to 15 degree interval but not the 15 to 20 degree interval. The frequency data were then normalized such that the interval with the highest number of species became 1 to remove any effect of total species richness. This gave us four sets of data based on a 2×2 factorial: sunbird and hummingbird polewardness and elevation. A logistic curve (eq. 1) was then fitted onto each of the four sets of data – the normalized species richness, *S*_*N*_, per interval vs. the midpoint of each interval – with variables *a* and *b* determining position and steepness of the curve respectively.

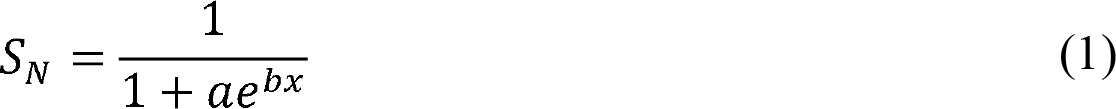

We then found the inflection point and the two points of the maximum magnitude of curvature (MMC points) for each curve. Inflection points indicate how well each family maintains species richness while MMC points give us the start and end of the decline in species richness. The functions and their key points characterize the shape of each family’s gradient.

## Results

Broadly, our results show that hummingbirds extend farther poleward and higher in elevation than do sunbirds. Hummingbirds extend from 62 degrees north to 56 degrees south and up to 5000 m in elevation (SI Table 1,2; Fig. 2,3). Sunbirds, on the other hand, extend only from 36 degrees north to 40 degrees south and up to 4880 m in elevation (SI Table 1,2; Fig. 2,3). Both families show the same general pattern of an initial increase in species richness followed by a decline moving poleward and to higher altitudes (Fig. 4a, b). In addition, hummingbirds maintain their species richness at higher elevations and more extreme latitudes than sunbirds. ECDF results confirm this difference in biogeography between hummingbirds and sunbirds with elevation constituting the greatest difference (SI Table 3, SI Fig. 1).

**Fig. 2.**
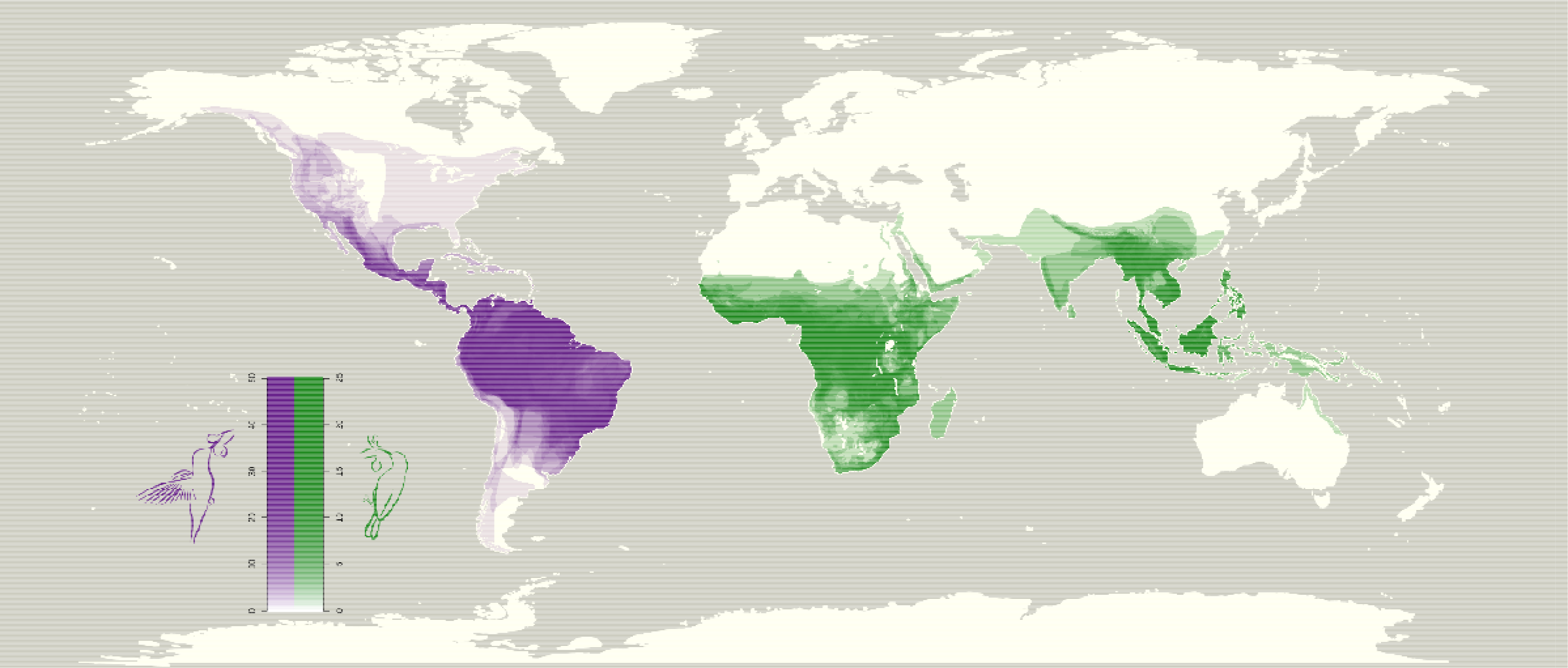
A map of species density of hummingbirds (purple) and sunbirds (green). Richer colors represent greater species density. Scales are chosen to reflect the difference in overall species richness of each taxon. Hummingbirds not only have higher species density but also extend farther.

**Fig. 3.**
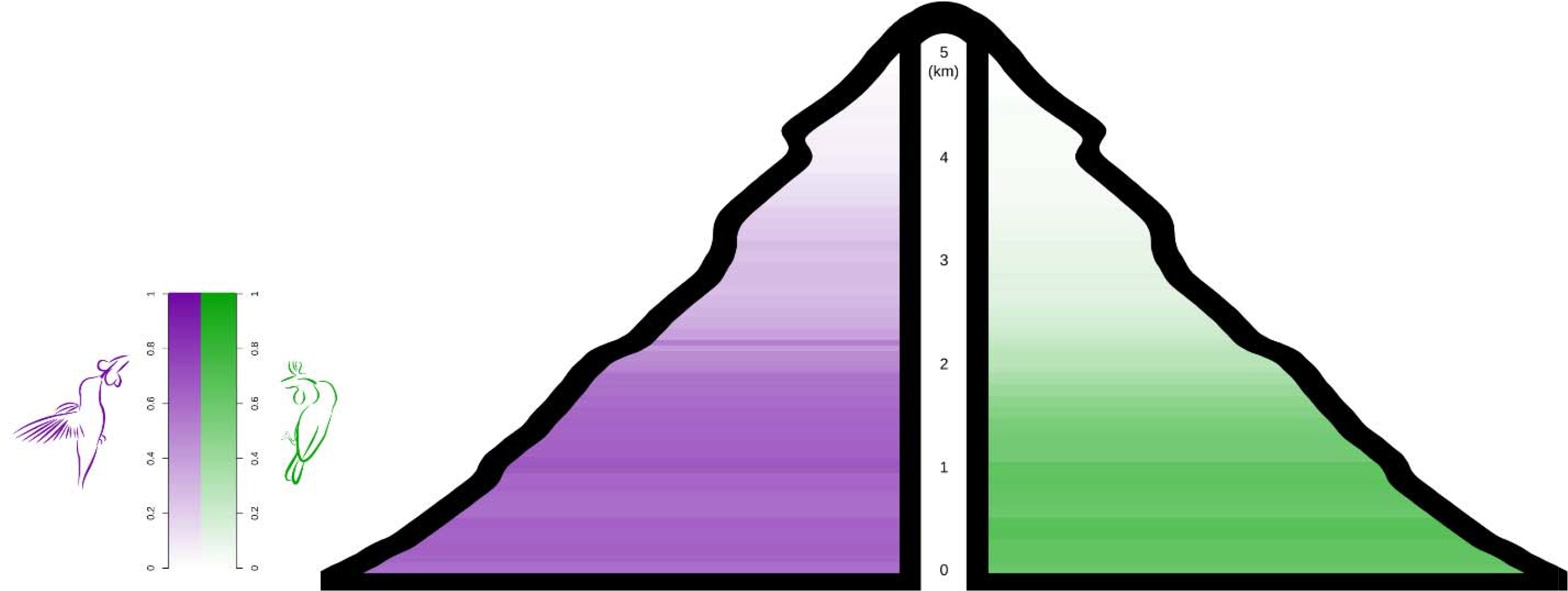
Changes in species density with elevation for hummingbirds (purple) and sunbirds (green). Though both clades extend to similar altitudes, hummingbirds maintain species richness at higher elevations as denoted by the richer colors.

**Fig. 4.**
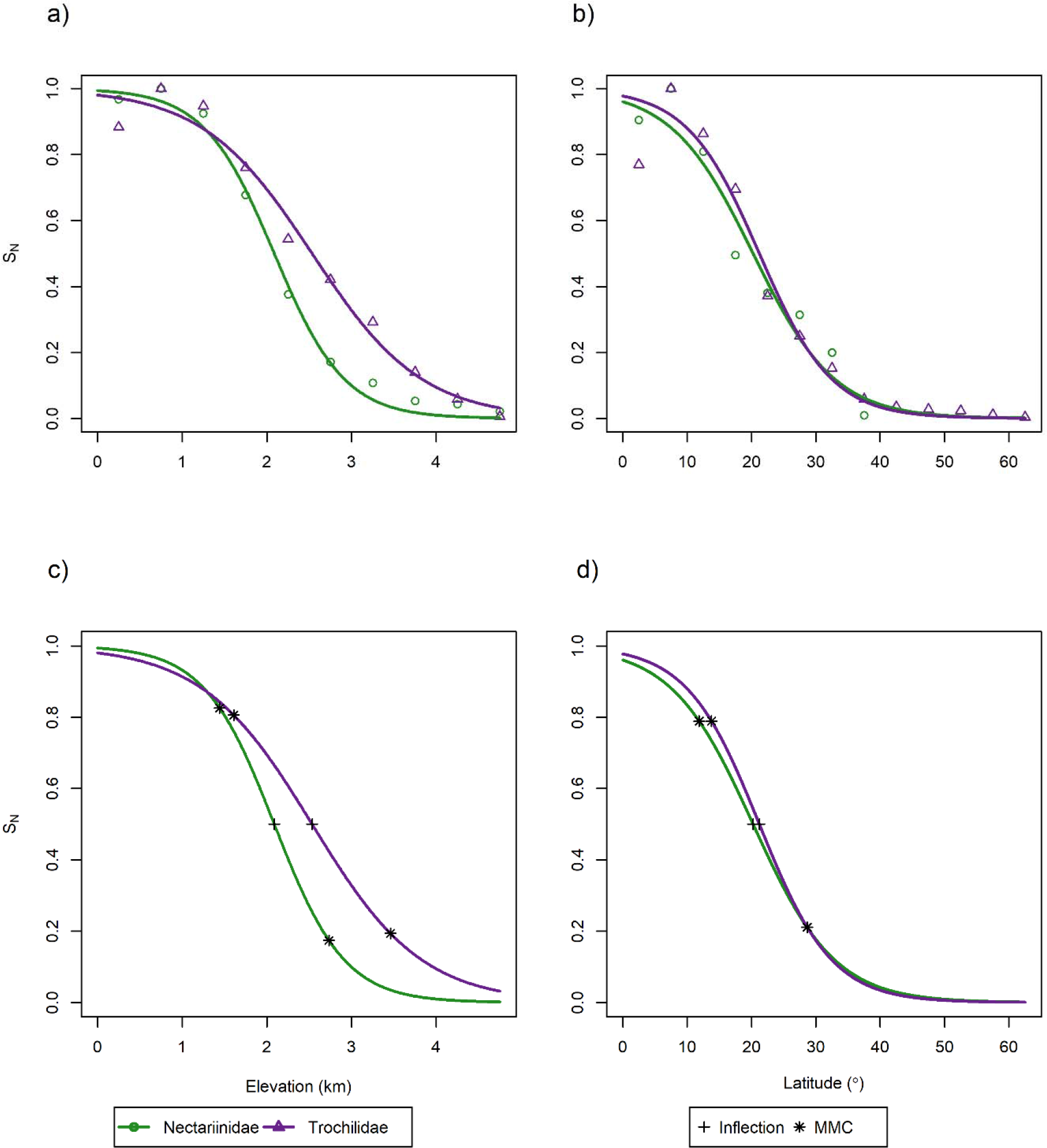
A plot of the normalized species richness *S*_*N*_ of hummingbirds and sunbirds, along with the fitted line, for elevation (a, c) and “polewardness” (b, d). Triangles and purple lines denote hummingbirds, and circles and green lines denote sunbirds. Hummingbirds maintain species richness at higher elevations and mid-latitudinal ranges and extend farther latitudinally than sunbirds. Inflection (cross) and MMC points (asterisks) also are shown (c, d). Inflection points come later in hummingbirds than sunbirds. With regard to elevation, hummingbird *S*_*N*_ and sunbird *S*_*N*_ start their decline at a similar spot but hummingbird *S_N_* declines more slowly. With latitude, sunbird *S*_*N*_ declines earlier than hummingbird *S*_*N*_

Both hummingbirds and sunbirds reach approximately the same maximum elevation, around 5000m (SI Table 1, SI Fig. 1a). Even though both hummingbirds and sunbirds extend to roughly the same elevation, hummingbirds have a higher normalized species richness at higher elevations compared to sunbirds. The inflection point for sunbirds occurs at 2087m and hummingbirds at 2533m (SI Table 4, Fig. 4c). Sunbirds and hummingbird species richness values both start to decline around the same elevation –1764 and 1898m respectively – but sunbirds plateau at a lower elevation compared to hummingbirds –2410m vs. 3458m respectively – indicating a more gradual decline in the normalized species richness of hummingbirds (SI Table 4, Fig. 4c).

Regarding latitude, hummingbirds occur farther from the equator than do sunbirds, 60-65 degrees vs. 35-40 degrees respectively (SI Table 2, SI Fig. 1b). Also, hummingbird normalized species richness is at its greatest divergence from sunbird normalized species richness at mid-latitudinal ranges. The hummingbirds’ inflection point is 22.14 degrees latitude versus 18.92 degrees for sunbirds (SI Table 4, Fig. 4d). Hummingbirds also begin their declines further from the equator than do sunbirds –14.99 and 9.44 degrees respectively. Both plateau around the same latitude – 29.29 vs. 28.39 degrees respectively (SI Table 4, Fig. 4d).

## Discussion

Sunbirds and hummingbirds are two convergent nectarivorous bird families with different evolutionary technologies. While hummingbirds are extremely specialized to nectar feeding, sunbirds vary, ranging from the highly specialized sugarbirds to the passerine-like flowerpeckers (10; 11). These differences in evolutionary technologies should reflect differences in the families’ distribution and biogeography. As one moves higher in elevation and towards the poles, hummingbirds maintain their species richness more than sunbirds. Though extending to roughly the same elevational maximum, normalized hummingbird species richness declines at a much slower rate than sunbirds. The same is true for latitude; in addition, hummingbirds extend into more extreme latitudes (farther north and south) than sunbirds. Clear from our results is that hummingbirds have a greater biogeographical extent than sunbirds, likely reflecting a superior key adaptation.

One potential hypothesis for the biogeographical differences of hummingbirds and sunbirds could be dispersal limitation. Firstly, there is a lack of suitable land below 40° S Secondly, Old World mountain ranges may form a barrier to sunbird dispersal as they primarily run along the east-west axis in contrast to New World mountain ranges which primarily run along a north-south axis. We reject this hypothesis on the grounds that hummingbirds are frequently found in montane habitats. Not only do hummingbirds maintain species richness at higher elevations as our study showed, they have higher species richness in the mountains of western North and South America compared to the flat-lying eastern regions and frequently undertake migrations in mountainous areas. Even if sunbirds were dispersal limited, hummingbirds are still more speciose than sunbirds even when taking latitudinal range into account. Of the 364 species, only 15 hummingbirds are found at latitudes where sunbirds are absent. Even if we assume that expansion into the northern latitudes led to the evolution of these 15 species, it still only accounts for approximately 4% of hummingbird species. The difference in species richness between the families cannot solely be due to dispersal limitation. Instead, we feel that the combined evidence of species richness and biogeography is highly suggestive of one or more key adaptations in hummingbirds.

Our spatial analyses cannot tell us what the key adaptations are, but we can speculate on what they may be. Though hummingbirds and highly specialized sunbirds show many similarities, they do differ in specific areas. Likely, the key adaptation deals with the differences in their foraging, specifically how they feed and how they fly. With feeding, one possibility for hummingbirds’ key adaptation may be their unique tongues. The tongues of hummingbirds have recently been shown to act as micropumps, a way of quickly and efficiently gathering nectar from flowers, in contrast to the previously assumed capillary action (13; 14). This unusual feeding method may allow hummingbirds to more efficiently gather nectar compared to sunbirds. Not enough is known about sunbird tongues, however, to see how the two taxa compare in nectar gathering abilities. Studies indicate that hummingbirds and sunbirds gather nectar at seemingly comparable rates suggesting that the amount gathered is not the key difference (31; 32; 33; 14 [personal calculation]). If the tongue is the key adaptation, it will be for the fact that micropumping requires no energy expenditure on the part of hummingbirds, which removes a cost, while sunbirds apparently intake nectar through suction, a potentially energetically expensive system (34; 35). More research needs to be done on the tongues of sunbirds to see how they compare with the tongues of hummingbirds.

Another possibility of the key adaptation that separates hummingbirds and sunbirds is hummingbirds’ ability to hover and fly in all directions (10). Adaptations for hovering include shortened arm bones, longer hand bones, a relatively fixed V-shaped arm position, a shallow ball-and-cup joint between the coracoid and sternum, a large sternum with a deep keel onto which large breast muscles – pectoralis and supracoracoideus – attach, and red blood cells and hemoglobin adapted for higher-oxygen affinity and carrying capability (36; 37; 39; 38). All these anatomical features are adaptations to stiff-winged flight and are seen to a lesser extreme within other bird families of the order Apodiformes (36; 37; 38). What truly differentiates the flight of hummingbirds is the axial rotation of the humerus and wrist bones during flight (38). Hummingbirds can create lift on the upstroke – in addition to the downstroke seen in all birds – due to wing inversion caused by axial rotation of the wrist (39). Wrist flexibility comes from changes in carpal structure and the deletion of key ligaments and is seen in birds outside of Apodiformes (40; 38; 41). Additional power for each downstroke and upstroke also comes from the axial rotation of humerus, driven by the pectoralis, supracoracoideus, and other muscles (42; 39; 38). The humerus can rotate up to 180º due to a unique humeroscapular joint (43; 36). In hummingbirds, the humeral head (condyle) is placed along the axis of the shaft instead of the terminal position, a feature unique to them (44; 45). Together, this suite of adaptations allows hummingbirds to hover effectively when foraging (46).

Other evolutionary technologies may also benefit hummingbirds in secondary ways. For example, hummingbirds sustain flight more efficiently at higher altitudes, likely due to their denser erythrocyte count, expanding their fundamental niche to higher elevations (47). We feel though that hovering remains the likeliest candidate for a hummingbird key adaptation. Many of the musculoskeletal changes are seen only in Apodiformes with the shifting of the condyle seen only in Trochilidae. Such efficient hovering is likely an evolutionarily implastic and ancestral trait that arose only once among Aves. Through this adaptation, hummingbirds have fundamentally changed the rules of their nectarivory; they exist as a new type of bauplan while sunbirds are still effectively a derived passerine (6; 48).

We speculate three possible reasons for the evolution of hovering. Firstly, hummingbirds can exploit the nectar of plants without perches, potentially opening a new resource for them. As other nectarivorous birds need to perch while feeding, flowers without perches may represent a relatively abundant and constant resource without competition from other bird species. Evolution of hovering in this scenario may be a virtuous cycle as hovering is more efficient at high nectar volumes which occur in the absence of competition (49). Secondly, hummingbirds may be better able to escape predation due to their unique flying abilities. With the ability to fly in all directions, hummingbirds may more easily avoid predators, a useful ability especially when feeding at a flower with blocked sightlines (50). Furthermore, the musculoskeletal changes in the hummingbirds are shown to make them extraordinarily agile (51). Finally, while hovering is energetically costly, it is also time efficient (52). Hovering birds spend less time gathering resources at flowers than birds which rely on perches. This means that hovering becomes more energetically efficient compared to perching when birds feed within clustered flower patches (53; 54).

There could be many reasons why hummingbirds developed their key adaptation. Hummingbirds underwent an expansive radiation during the uplift of the Andes beginning around 10 mya (55). Living in such rapidly changing conditions could have necessitated the evolution of a more efficient foraging system. As mentioned earlier, greater oxygen capacity is beneficial to both hovering and living in low oxygen conditions. There is also the possibility that the rise of the Andes freed up niche space that would have otherwise been taken up by a competing family like hawkmoths (Sphingidae), a sort of ecological and evolutionary constraint (5). These factors, along with hummingbirds’ evolutionary history, may combine to lead to the evolution of hovering (46). Furthermore, sunbirds may face their own internal constraints, genetic or otherwise, preventing them from evolving a key adaptation (56). Whatever the case may be, our results suggest the evolution of hovering (or some other adaptation) allowed hummingbirds to more efficiently take advantage of a resource and expand their fundamental niche.

The real test of evolutionary technologies would come from seeing what happens when the two clades meet. Deliberately shifting species across the globe would obviously be unethical but previous or accidental species invasions may offer such a test. For example, European Lumbricid earthworms have colonized parts of North America that are farther north than their American counterparts (57). Both sets of earthworms are ecological equivalents and have convergent features to fill the role of soil turners. The invasive European earthworms, though, are known to tolerate environmental stress through protective cocoons during times of drought and high glucose and glycogen content in cells to prevent freezing during winter (58; 59). These adaptations may have allowed European earthworms to colonize the colder climes of Canada and expand their range beyond the North American species.

Through biogeographic analysis, we show that hummingbirds inhabit more hostile climes than sunbirds, likely due to the possession of a superior evolutionary technology. Going forward, biogeographic comparison between clades may reveal itself to be a powerful tool to reveal differences in evolutionary technologies and illuminate the interaction between adaptation and environment.

## Supporting information

Additional Results, Figures, and Methods

## Acknowledgements

Abdel Halloway wishes to thank the NSF for funding his graduate studies. This material is based upon work supported by the National Science Foundation Graduate Research Fellowship under Grant Nos. DGE-0907994 and DGE-1444315. Any opinion, findings, and conclusions or recommendations expressed in this material are those of the authors(s) and do not necessarily reflect the views of the National Science Foundation.

