## Additional Results, Figures, and Methods for "Using Biogeography to Assess Key Adaptation Strength in Two Bird Families"

SI Figure 1. A plot of four select ECDFs used to compare hummingbird and sunbird distribution. Purple lines indicate hummingbirds and green lines indicate sunbirds. Hummingbird ECDFs almost entirely lie below sunbird ECDFs with the greatest difference occurring with elevation measures.

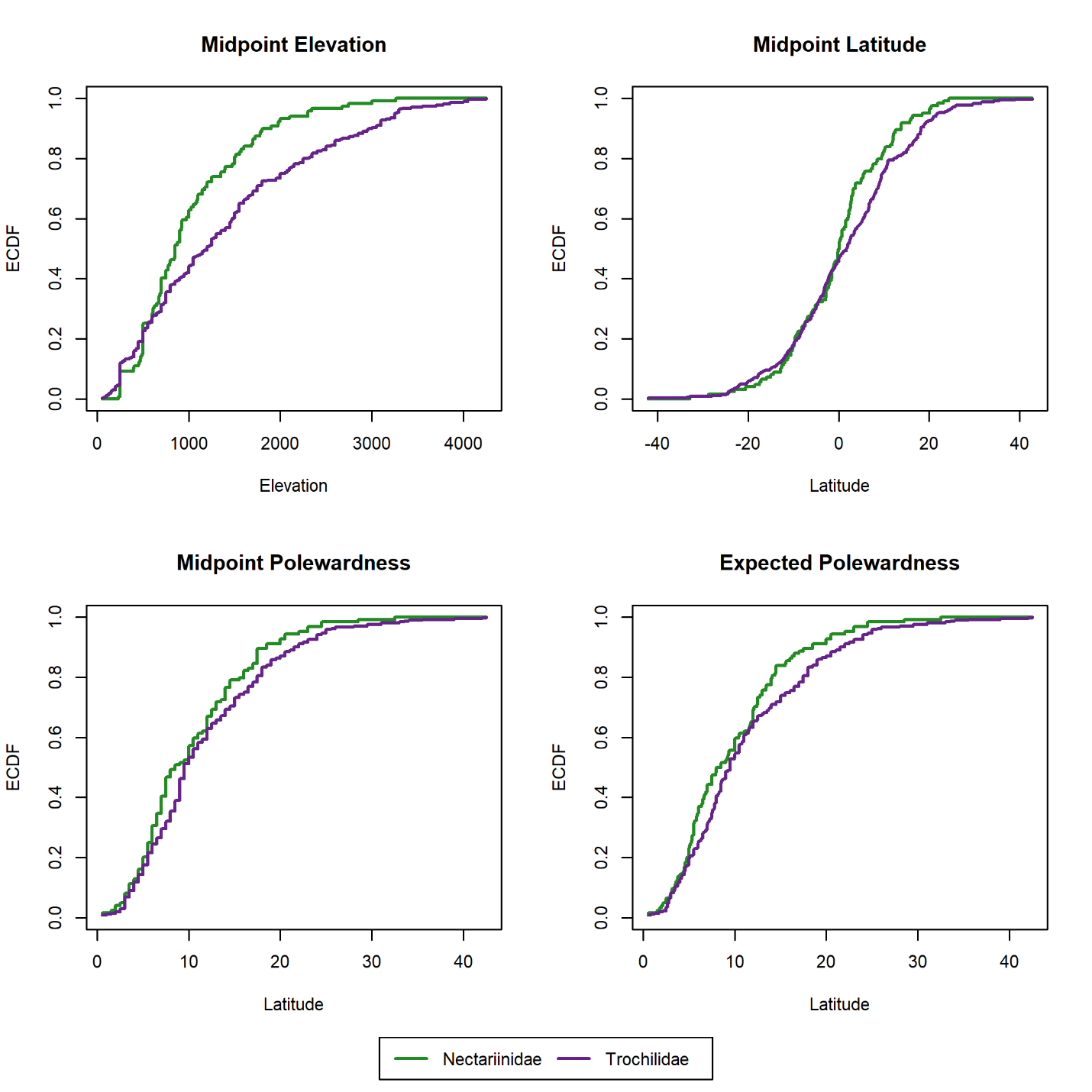

SI Table 1: Normalized species richness per elevation interval of sunbirds and hummingbirds

| **Elevation**  **(meters)** | **Sunbirds** | **Hummingbirds** |
| --- | --- | --- |
| [ 0 , 500 ] | 0.967741935 | 0.883040936 |
| ( 500 , 1000] | 1 | 1 |
| (1000 , 1500] | 0.924731183 | 0.947368421 |
| (1500 , 2000] | 0.677419355 | 0.760233918 |
| (2000 , 2500] | 0.376344086 | 0.543859649 |
| (2500 , 3000] | 0.172043011 | 0.421052632 |
| (3000 , 3500] | 0.107526882 | 0.292397661 |
| (3500 , 4000] | 0.053763441 | 0.140350877 |
| (4000 , 4500] | 0.043010753 | 0.058479532 |
| (4500 , 5000] | 0.021505376 | 0.005847953 |

SI Table 2: Normalized species richness per latitudinal interval of sunbirds and hummingbirds

| **Latitude**  **(degrees)** | **Sunbirds** | **Hummingbirds** |
| --- | --- | --- |
| [ 0 , 5 ] | 0.904761905 | 0.76953125 |
| ( 5 , 10 ] | 1 | 1 |
| ( 10 , 15 ] | 0.80952381 | 0.86328125 |
| ( 15 , 20 ] | 0.495238095 | 0.6953125 |
| ( 20 , 25 ] | 0.380952381 | 0.37109375 |
| ( 25 , 30 ] | 0.314285714 | 0.25 |
| ( 30 , 35 ] | 0.2 | 0.15234375 |
| ( 35 , 40 ] | 0.00952381 | 0.05859375 |
| ( 40 , 45 ] | 0 | 0.03515625 |
| ( 45 , 50 ] | 0 | 0.02734375 |
| ( 50 , 55 ] | 0 | 0.0234375 |
| ( 55 , 60 ] | 0 | 0.01171875 |
| ( 60 , 65 ] | 0 | 0.00390625 |

SI Table 3: The results of the ECDF comparisons (MDE tests) between families. *Italics* indicate the centrality based ECDFs. **KS** stands for Kolmogorov-Smirnov and **AD** stands for Anderson-Darling. NS indicates that the result was not significant.

| **ECDF Type** | **Hbird Num** | **Sbird Num** | **KS**  **D-statistic** | **KS**  **Significance** | **Standardized AD Criterion** | **AD**  **Significance** |
| --- | --- | --- | --- | --- | --- | --- |
| Minimum  Elevation | 309 | 119 | 0.331783199 | p<0.001 | 31.482 | p<0.001 |
| Maximum  Elevation | 309 | 119 | 0.163280846 | p<0.05 | 3.2666 | p<0.01 |
| *Midpoint*  *Elevation* | 309 | 119 | 0.214598461 | p<0.001 | 8.685 | p<0.001 |
| Minimum  Latitude | 365 | 124 | 0.234423332 | p<0.001 | 10.207 | p<0.001 |
| Maximum  Latitude | 365 | 124 | 0.069443217 | NS | 0.49404 | NS |
| *Midpoint*  *Latitude* | 365 | 124 | 0.156407424 | p<0.05 | 1.7652 | p<0.05 |
| Minimum  Polewardness | 365 | 124 | 0.242222713 | p<0.001 | 14.455 | p<0.001 |
| Maximum  Polewardness | 365 | 124 | 0.073486522 | NS | 0.71488 | p<0.1 |
| *Midpoint*  *Polewardness* | 365 | 124 | 0.14719399 | p<0.05 | 1.4571 | p<0.05 |
| *Expected*  *Polewardness* | 365 | 124 | 0.13121962 | p<0.05 | 1.8851 | p<0.05 |

SI Table 4: The values of $a$ and $b$, the inflection point, and the MMC points for each of the models along with their significance. RSS is the residual sum of squares for each model, and RSE is the residual standard error. It should be noted that all residuals fall between 0 and 1 which may exaggerate the RSS.

| **Model** | **a** | **b** | **Inflection** | **MMC #1** | **MMC #2** | **RSS** | **RSE** |
| --- | --- | --- | --- | --- | --- | --- | --- |
| NectarElev | 0.00655^d^ | 2.40963^a^ | 2.087005 | 1.441369 | 2.73264 | 0.010254 | 0.035802 |
| TrochElev | 0.02024^c^ | 1.53943^a^ | 2.533536 | 1.608667 | 3.458399 | 0.022979 | 0.053594 |
| NectarLat | 0.04133^c^ | 0.15735^a^ | 20.24934 | 11.872 | 28.62667 | 0.04463 | 0.063696 |
| TrochLat | 0.02318 | 0.17739^a^ | 21.22319 | 13.79037 | 28.656 | 0.054938 | 0.070671 |

a: p<0.001, b: p<0.01, c: p<0.05, d: p<0.1

**Additional Methods**

*More on Latitudinal and Elevational Gradients*

Besides the geographic and environmental reasons that make latitudinal and elevational gradients useful for this study, there are other methodological features that make them particularly useful for this analysis. Spatial comparisons over a wide geographic range are better able to indicate the presence of key adaptations, as they cover many environmental variables – reducing the chance of a false negative – but are not inherently correlated to the adaptation in question – preventing false positives. Additionally, elevational comparisons offer a stronger comparison between families. This is because a random distribution of the range sizes and locations within a bounded geographical space of a group of species will generate a hump-shaped gradient broadly resembling (though not identical to) latitudinal gradients (1). Elevational gradients occur in bounded geographical space but present a skewed distribution indicating the lack of random processes. Therefore, any difference between the families is likely an effect of their natural histories.

*Expected Polewardness*

The measure of expected polewardness was calculated with the assumption that abundance was stacked for species whose ranges crossed the equator. We assumed a uniform distribution of abundance across each species range We calculated the expected value of this stacked uniform distribution seen in eq. (1) where $min$ and $max$ stand for the absolute value of minimum and maximum latitude respectively before folding.

|  | $\frac{1}{2}*\frac{min^{2}+max^{2}}{\left( min+max \right)}$ | (1) |
| --- | --- | --- |

1. Willig MR, Lyons SK. An analytical model of latitudinal gradients of species richness with an empirical test for marsupials and bats in the New World. Oikos. 1998 Feb 1:93-8.
